# Concordance between the fast and efficient mixed-effects algorithm (FEMA) and conventional mixed-effects modeling for whole-brain connectivity in longitudinal chronic pain

**DOI:** 10.64898/2026.09.20.753017

**Authors:** Natalie McLain, Chelsea Kaplan, Steven Harte, Andrew Schrepf

## Abstract

Researchers increasingly have access to neuroimaging data with repeated measures within subjects. Linear mixed-effects modeling offers a valuable way to analyze such data, but its application remains limited in the neuroimaging literature due to high computational cost and a lack of mainstream analysis packages. Here we benchmark a recently published algorithm, the Fast and Efficient Mixed-Effects Algorithm (FEMA), against a conventional implementation (MATLAB’s fitlme, restricted maximum likelihood) across 94,830 functional connectivity edges. In contrast to the Adolescent Brain Cognitive Development (ABCD) Study Dataset used in the development and testing of FEMA, the dataset used in the current analysis has a smaller sample size (n = 378) and comprises individuals with chronic pelvic pain scanned at up to four visits over three years. Despite these differences, the two implementations produced highly concordant results: fixed-effect estimates and test statistics correlated at r ≥ 0.999 (Lin’s concordance correlation coefficient ≥ 0.999), and the two approaches reached the same statistical conclusion for 99.98% of edges. FEMA completed each analysis roughly 20 times faster. Agreement was weaker for the estimated variance components, where FEMA attributed systematically less variance to the participant random effect, yielding a slightly lower intraclass correlation in approximately 82% of edges. We further show that most residual disagreement reflected FEMA’s default variance-grid resolution rather than the estimator itself, and could be eliminated at negligible computational cost. These findings support the use of FEMA for connectome-wide analysis of repeated-measures neuroimaging data.

## 1 Introduction

Identifying robust relationships between brain function and chronic pain remains a central challenge in neuroimaging. While cross-sectional analyses have provided valuable insight about group differences between individuals with chronic pain and healthy controls, these approaches often obscure substantial within-subject variability. In addition to being biologically meaningful, this variability impacts the reliability of functional magnetic resonance imaging (fMRI) measures (Flournoy et al., 2024; Nakuci et al., 2023; Noble et al., 2019, 2021; Tozzi et al., 2020). Indeed, functional connectivity specifically has shown overall poor reliability, with the largest meta-analysis of test-retest reliability in functional connectivity studies showing a field-wide average of approximately 0.3 (Noble et al., 2019). These concerns are especially relevant in the context of longitudinal cohort studies that image tens of thousands of individuals (Fiúza-Fernandes et al., 2025; Henn et al., 2023; Kim et al., 2021; Tanasescu et al., 2016).

Increasingly, neuroimaging resources such as the UK Biobank and the Adolescent Brain Cognitive Development (ABCD) study include repeated measurements and nested data structures alongside rich clinical phenotyping (Hsu et al., 2025; Kaplan et al., 2022; McQueenie et al., 2021). This presents opportunities to examine symptom-brain relationships across and within individuals over time. However, conventional analytic approaches often treat repeated observations as independent or rely on summary measures that discard within-subject information, limiting sensitivity to clinically relevant effects (Madhyastha et al., 2018).

One viable method for addressing this issue is linear mixed-effects modeling (LMEs) paired with repeated measures designs. By explicitly modeling subject-level dependence, LMEs can account for repeated measures, hierarchical data structures, and unbalanced designs (Chen et al., 2013). However, the substantial advantages offered by LMEs are rarely employed in neuroimaging analyses because of intensive computational demands and a lack of mainstream toolboxes (Chen et al., 2013; Madhyastha et al., 2018; Parekh et al., 2024; Russman Block et al., 2023). This is especially true in applications such as whole-brain analyses which can involve tens of thousands of models.

Recently, the Fast and Efficient Mixed-Effects Algorithm (FEMA) was introduced to address these limitations (Parekh et al., 2024). FEMA enables mass-univariate LME analyses by dramatically reducing computation time through a combination of method-of-moments variance estimation and generalized least squares, while preserving the ability to model complex longitudinal and hierarchical data structures (Parekh et al., 2024). In its original validation, FEMA demonstrated strong agreement with conventional likelihood-based mixed-effects solvers, alongside orders-of-magnitude improvements in computational efficiency. FEMA has since been recommended for longitudinal analyses within the ABCD study (Hawes et al., 2025), the dataset it was originally validated and developed in.

Although FEMA shows strong promise, it has seen limited application outside of the dataset used in its original development. Specifically, it has yet to be demonstrated whether FEMA produces results that are concordant with conventional mixed-effects approaches when applied to repeated-measures functional connectivity data in smaller data sets of chronic pain patients, where effect sizes are often modest and within-subject variability is substantial.

Here, we apply FEMA to a longitudinal dataset of individuals with chronic pelvic pain and compare its performance against a conventional linear mixed-effects implementation using MATLAB’s fitlme function. We focus on whole-brain, node-wise functional connectivity and examine two complementary dimensions of the pain experience: pain reported at the time of scanning and recalled average pain over the preceding week. Our primary goal is not to introduce new neurobiological claims, but rather to evaluate the extent to which FEMA reproduces established mixed-effects results (McLain et al., 2026) with a smaller number of subjects in a dataset and modeling context that were not part of its original validation. Specifically, we assess concordance between methods in terms of identified pain-related connectivity patterns, fixed-effect estimates, and statistical inference, while also quantifying differences in computational efficiency.

By benchmarking FEMA in a longitudinal chronic pain dataset and in the context of connectome-wide functional connectivity, this work aims to inform the broader adoption of mixed-effects modeling in neuroimaging.

## 2 Materials and Methods

This work is based, in-part, on an earlier publication (McLain et al., 2026). The current approach allowed us to compare FEMA against a known set of results in a dataset/set of models not previously tested or used in FEMA’s development. Methods for the fitlme approach were described in the earlier publication (McLain et al., 2026). Here we briefly describe the previous methods as well as methods that were added to or differ from the original approach.

### 2.1 Participants

Data were drawn from the Multidisciplinary Approach to the Study of Pelvic Pain (MAPP) Symptom Patterns Study (Clemens et al., 2020) where participants with urologic chronic pelvic pain scanned repeatedly (2-4 visits) over 36 months. Of the 492 chronic pain patients, the final number of subject included in this analysis was 378.

Filtering criteria can be found in section 2.6 Data Filtering. At each visit, resting-state fMRI was acquired during two naturalistic bladder states: a “fuller bladder” scan following controlled water intake and an “empty bladder” scan after voiding. Participants reported pain at the time of each scan and recalled average pain over the preceding week.

#### 2.1.1 Ethics Statement

All study protocols were approved by the Institutional Review Boards at the six study sites. All study protocols were followed according to the Declaration of Helsinki. All participants provided informed consent for participating in the study.

#### 2.1.2 Inclusion and Exclusion Criteria

Inclusion and exclusion criteria have been previously published (Clemens et al., 2020). Briefly, UCPPS inclusion criteria were 1) UCPPS symptoms present for a majority of the time during most recent 3 months; 2) age ≥18 years; and 3) response ≥1 on the bladder/prostate or pelvic pain/pressure/discomfort scale during past 2 weeks. Exclusion criteria include symptomatic urethral stricture, neurological disease or disorder affecting the bladder, bladder fistula, a history of cystitis caused by tuberculosis, radiation therapy or chemotherapy, prior augmentation cystoplasty or cystectomy, active autoimmune or infectious disorder, history of pelvic cancer, current major psychiatric disorder, severe cardiac, pulmonary, renal, or hepatic disease, unilateral orchialgia (without pelvic symptoms), and prior prostate procedures (transurethral microwave thermotherapy (TUMT), transurethral needle ablation (TUNA), balloon dilation, prostate cryo-surgery, or laser procedure).

### 2.2 MRI Protocol

The SPS MRI acquisition protocol has been described previously (Clemens et al., 2020; Mawla et al., 2020). Briefly, participants emptied their bladders and then consumed 350cc of water. After approximately 40 minutes, a resting state fMRI (rs-fMRI) scan labelled “fuller bladder” (rs-FB) was performed (10 min acquisition). Following rs-FB, participants exited the scanner and emptied their bladders. Returning to the scanner, an “empty bladder” resting state (rs-EB) scan was performed (10 min acquisition), followed by a T1-weighted structural scan.

#### 2.2.1 MRI Acquisition

The Neuroimaging Core of the G. Oppenheimer Center for Neurobiology of Stress and Resilience (CNSR) at UCLA operated as the neuroimaging data coordinating hub for the MAPP Research Network. Scanning took place at six data collection sites: Northwestern University (NU), Chicago, Illinois; University of California/University of Southern California, Los Angeles (UCLA/USC); University of Iowa (UI), Iowa City; University of Michigan (UM), Ann Arbor; University of Washington (UW), Seattle; and Washington University (WashU), St. Louis, Missouri. The full scanning parameters and quality control procedures for multi-site rs-fMRI and T1 imaging have been described previously (Mawla et al., 2020).

Here we briefly describe the nominal rs-fMRI and T1 parameters: rs-fMRI scans were acquired with a single shot echo planar imaging (EPI) pulse sequences with conventional rectangular Cartesian sampling. Basic pulse sequence parameters were as follows TR = 2000 ms, TE = 30 ms, Flip angle = 77°, FoV = 220 mm × 220 mm, Resolution = 64 × 64, Phase encode direction = A > P, Slice thickness = 4.0 mm, Slice gap = 0.5 mm, Slice acquisition = ascending (not interleaved), Slices per volume = 34-40 to cover entire brain, Phased array acceleration factor = 2, Bandwidth = maximum to accommodate resolution specifications, Orientation = axial-oblique parallel to the line between the anterior and posterior commissures, Number of volumes = 300 (10 min acquisition).

The MP-RAGE pulse sequence was used for high-resolution T1-weighted 3D volume imaging. The equivalent pulse sequence on a GE scanner was the 3D FSPGR IR. Basic parameters were as follows: TR = 2300 ms, TE = 2.98 ms, TI = 900 ms, Flip angle = GE: 11°, Siemens: 9°, Philips: 9°, FoV = 256 mm × 256 mm, Resolution = 256 × 256, Slices per volume = 240 (or maximum available while maintaining all other parameters), Slice thickness = 1 mm, Inversion = Slice Selective, parallel imaging acceleration factor = 2, Phase encode direction = left-right and superior-inferior, Orientation = axial-oblique parallel to the line between the anterior and posterior commissures.

#### 2.2.2 rs-fMRI and T1 Preprocessing

rs-fMRI and T1 preprocessing were performed at the Chronic Pain and Fatigue Research Center, University of Michigan, Ann Arbor, MI. Results included in this manuscript come from preprocessing performed using fMRIPrep 20.2.0 (Esteban et al., 2019); RRID:SCR 016216, which is based on Nipype 1.5.1 (Esteban et al., 2022; Gorgolewski et al., 2011); RRID:SCR 002502.

#### 2.2.3 T1 Preprocessing

The T1-weighted (T1w) image underwent intensity correction using N4BiasFieldCorrection (ANTs 2.2.0) (Tustison et al., 2010) and skull-stripping using antsBrainExtraction.sh (ANTs 2.2.0). Surface reconstruction was performed with recon-all (FreeSurfer 6.0.1) (Dale et al., 1999), with additional refinement of the brainmask (Klein et al., 2017). Nonlinear registration and spatial normalization to the 2009c ICBM152 template was performed with antsRegistration (ANTs 2.2.0) (Avants et al., 2008). Tissue segmentation of cerebrospinal fluid (CSF), white-matter (WM), and gray-matter (GM) was performed on the brain-extracted T1w using fast (FSL 5.0.9).

#### 2.2.4 Functional Data Preprocessing

For each of the 2 BOLD runs per subject (rs-EB and rs-FB scans at each visit), the following preprocessing was performed. Reference images were co-registered with 9 degrees-of-freedom to the T1w reference using boundary-based registration (bbregister, FreeSurfer) (Greve & Fischl, 2009). Head-motion realignment was performed using mcflirt (FSL 5.0.9) (Jenkinson et al., 2002). Images were warped to MNI152NLin2009cAsym standard space and resampled to 2 × 2 × 2 mm voxel dimension to allow for cross-subject comparison. Framewise Displacement (FD) was calculated using Nipype (Power et al., 2014). Six physiological regressors were extracted for principal component-based noise correction based on anatomical CSF and WM masks computed in native space (aCompCor) (Behzadi et al., 2007). Following the fMRIprep minimally preprocessed pipeline, the preproc.nii images were skull-stripped using the subject-specific brain mask generated by fMRIPrep in MNI152NLin2009cAsym space, which was slightly dilated using fslmaths to ensure full cortical coverage. Based on recent recommendations (Lindquist et al., 2019), six head-motion parameters from mcflirt, six aCompCor regressors, and high-pass temporal filtering (0.01 Hz) were simultaneously regressed out of the BOLD time series using AFNI’s 3dTproject function. Finally, 3DBlurToFWHM was used to estimate smoothness of each image followed by iterative smoothing until the images reached a target smoothness of 6 mm FWHM.

### 2.3 Motion Assessment

Two motion criteria were applied to exclude high-motion scans: maximum framewise displacement (FD) greater than 3 mm and mean FD greater than 0.5 mm. Out of 3,108 rs-FB and rs-EB runs, 10 subjects and 681 scans were excluded based on motion filtering.

### 2.4 Connectivity Matrix Calculation

Connectivity matrices were calculated using the Schaefer+Brainnetome Subcortex parcellation scheme, which consists of 436 nodes: 400 cortical nodes derived from the Schaefer atlas (Schaefer et al., 2018) and 36 subcortical nodes derived from the Brainnetome atlas (Fan et al., 2016). Each node is designated to one of 8 networks: Visual (Vis), Somatosensory/Motor (SomMot), Limbic, Dorsal Attention (DorsAttn), Salience/Ventral Attention (SalVentAttn), Executive Control (Cont), and Default Mode (Default) from Schaefer and Subcortex for nodes from the Brainnetome atlas. Nodes have associated MNI-space centroids for visualization.

Post-processed, four-dimensional, rs-FB and rs-EB images were entered into nilearn. Functions under nilearn.connectome (e.g., ‘ConnectivityMeasure’) was used to compute Fisher z-transformed bivariate correlation (Pearson’s r) matrices of the 436x436 density matrix. In the resulting density matrix, each index represents the measure of functional connectivity, f, calculated between a pair of nodes. Each matrix contained f values for a total of 94,830 unique edges.

### 2.5 Pain Assessment

Two measures of pain were used in this analysis. Recalled pain was the average self-reported genitourinary pain over the week prior to the scanning visit as assessed with responses on the Genitourinary Pain Index (GUPI), question number four: “Which number best describes your AVERAGE pain or discomfort on the days you had it, over the last week?” Possible scores range from 0-10 (Clemens et al., 2009). Pain at time of scan for both empty and full scans was calculated as an average of pain intensity ratings (0-10) measured immediately before and immediately after each scan: “Several times during the session we will ask you to rate pain and urgency to urinate. Please rate pain from 0 (no pain) to 10 (worst pain imaginable) and urgency from 0 (no urge to urinate) to 10 (worst urge imaginable).”

### 2.6 Data Filtering

Analyses were restricted to participants meeting motion criteria (worst framewise displacement <3 mm; mean <0.5 mm, see motion assessment section above). 10 participants (681 scans) were dropped as a result of this filtering step. This left a total of 482 participants (2427 scans).

In a repeated measure, mixed effects model, the variance associated with the slope of a participant with only one data point is large and, as a result, their data will have little leverage on the group-level slope estimate. As a precautionary measure and to avoid inflating the degrees of freedom, we included only a subset of the original sample: individuals who completed at least two of the four study visits. After excluding the 104 remaining individuals who completed only one visit, 378 participants remained (2234 observations).

A complete-case filtering procedure was applied globally to remove scans with missing imaging values or missing model covariates, ensuring consistent sample sizes across all edges. 6 scans were dropped as a result of this complete-case filtering, but the number of subjects stayed the same. Site effects were harmonized using ComBat with biological covariates preserved. The final dataset comprised 378 participants (2228 scans). This final dataset was used in both the fitlme and FEMA implementations (described below), so both approaches were using identical data.

### 2.7 Statistical Approach

Two linear mixed-effects models were fit separately for pain at the time of scan and recalled pain, adjusting for age, sex, scan type, and head motion, with a random intercept for participant.

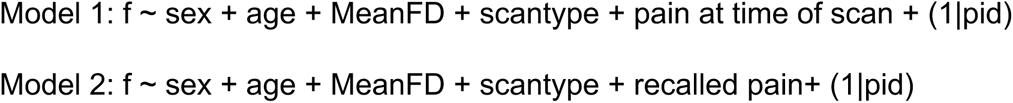

We ran Models 1 and 2 using both FEMA and a more basic set of scripts that relied on the fitlme function in MATLAB.

The fitlme approach relies on the base linear mixed effects modeling function in Matlab which uses restricted maximum likelihood (REML) for variance estimation. Fitlme estimates the full random-effects covariance matrix via a Cholesky parameterization. The maximum number of optimization iterations was set to the default of 1000. Additionally, all model outputs from the fitlme approach were assessed to confirm convergence. Of the 94,830 models per pain measure, none failed to converge; 183 (pain at time of scan) and 164 (recalled pain) returned a “local minimum possible” exit status, indicating that the gradient tolerance was satisfied but optimality could not be certified. None of the “local minimum possible” edges survived FDR correction (described below). Degrees of freedom for fixed effects were approximated using the Satterthwaite method, which has been shown to produce consistent Type I error rates and be more conservative than other common approaches for approximating p-values in linear mixed-effects models (Luke, 2017).

FEMA implementation was based on the same models, Models 1 and 2. A random intercept for participant ID was specified, with the family and individual identifiers set to the participant level. Model fitting was performed using the FEMA_fit function from the CMIG Tools FEMA implementation which uses generalized least squares (GLS) to estimate mixed-effects parameters, and a method-of-moments (MoM) approach to estimate the variance components for random effects. Unlike iterative likelihood-based optimizers used for ML/REML in standard mixed-effects solvers, FEMA’s default MoM + GLS approach does not rely on iterative likelihood maximization (ML/REML) (Parekh et al., 2024). Further details on the FEMA function and its modeling approach are comprehensively outlined in the original publication (Parekh et al., 2024) and associated materials on the authors’ github repository (https://github.com/cmig-research-group/cmig_tools).

For both implementations, fitlme and FEMA, p-values for the fixed effects of pain at time of scan and recalled pain were false discovery rate (FDR) corrected across all edges (94,830) using the Benjamini-Hochberg procedure (via the mafdr function in MATLAB), with a significance threshold of q < 0.01.

### 2.8 Assessing Concordance

Agreement between the two implementations was assessed at two levels: across the complete set of 94,830 edgewise models, independent of statistical significance, and again examining only the set of significant edges from each approach.

For the complete set of edges, we compared the standardized fixed effect of pain, its standard error, the corresponding test statistic, the signed −log10 p value, and the estimated variance components. Test statistics were computed from each implementation’s own standard error (Satterthwaite-adjusted t for fitlme, Wald z for FEMA). Agreement was quantified using both the Pearson correlation, a measure of precision, and Lin’s concordance correlation coefficient (CCC) (L. I.-K. Lin, 1989), which factors in both accuracy and precision coefficients (L. Lin et al., 2002; L. I.-K. Lin, 1989). Differences between implementations (FEMA minus fitlme) were further summarized as the median, interquartile range, and 95% limits of agreement (mean difference ± 1.96 SD), and displayed using Bland-Altman plots, in which the difference between methods was plotted against their mean (Bland & Altman, 1986; Giavarina, 2015; Mansournia et al., 2021).

To examine only the significant edges, we quantified set overlap with the Dice and Jaccard coefficients. Significant effects were then visualized using custom MATLAB scripts to create glass-brain representations, with edge color indicating effect direction and magnitude, and node size reflecting the number of significant connections per region.

FEMA discretizes the estimated random-effect variance into a grid and computes a single generalized least squares solution for all outcomes falling within the same grid cell, rather than solving separately for each outcome. The default resolution is set to 20 bins, but can be adjusted by the user. To determine whether this approximation contributed to any differences observed between implementations, the FEMA analyses were repeated at grid resolutions of 100, 1000, and 0 bins (one solution per edge, no binning). All other settings were held constant, and each was compared against the original/fitlme reference.

### 2.9 Use of AI

The original analysis pipeline was designed and written by the authors. Claude (Anthropic, model “claude-opus-5”) was used as an assistant throughout to review the code and identify bugs, parameterize an older version of the code to be more flexible for future applications, to add derived summary statistics during preparation of the associated manuscript, and to clean and package the repository for publication. All AI generated/edited code was compared against a known set of results. Specific applications of AI for each code file can be found in the repository README.

## 3 Results

### 3.1 Demographic Information

378 subjects with a minimum of two visits were entered into this analysis (240 female / 138 male). At baseline, mean age (mean±SD) across this study population was 44.7±15.7 years and mean symptom duration was 12.1±11.8 years. Mean age was 42.6±15.6 for females and 48.5±15.4 for males. Mean duration of symptoms was 10.8±10.4 years for males and 12.8±12.5 years for females. Further demographic data is summarized in Table 1.

**Table 1.**
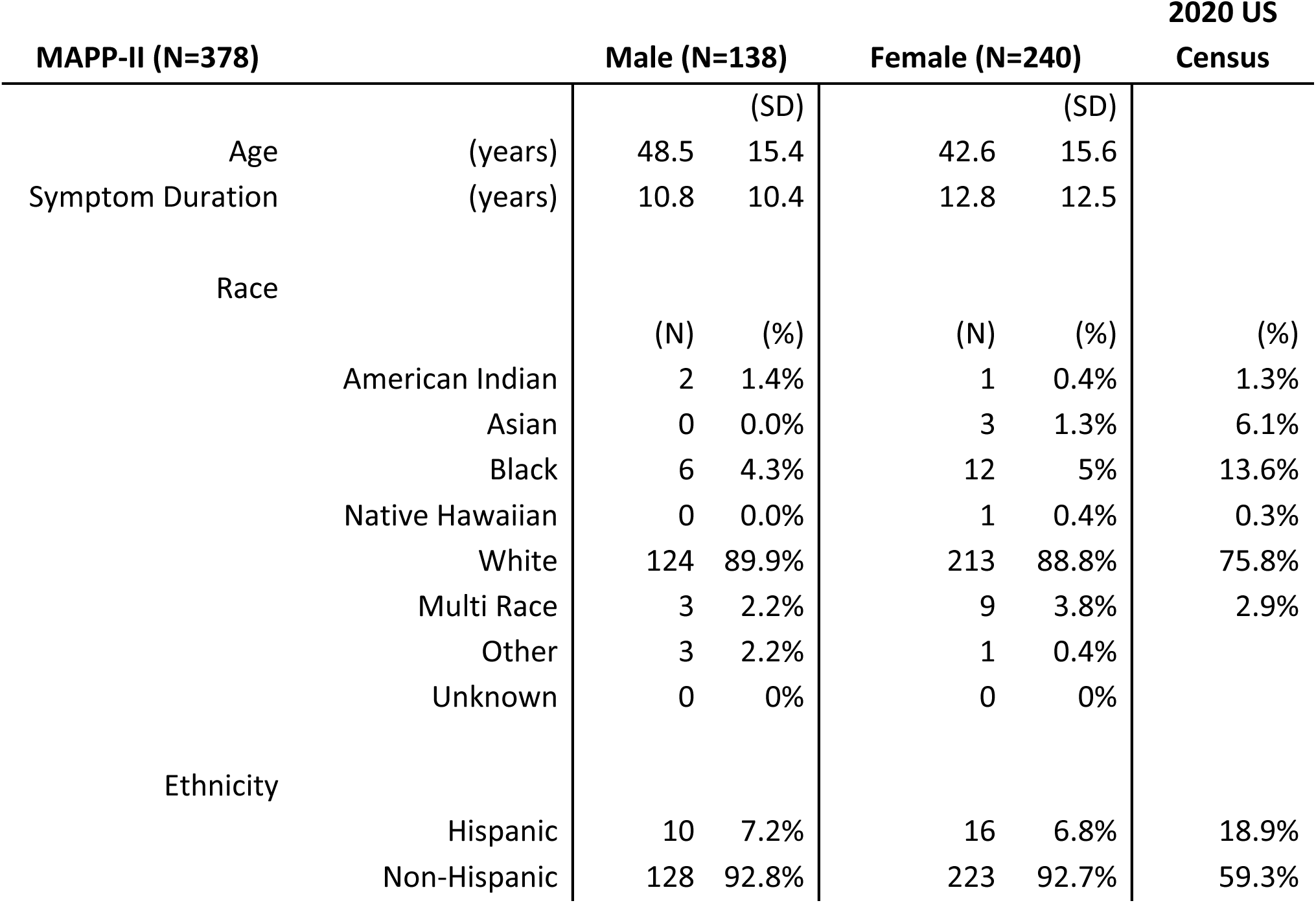
The demographic information for the 378 subjects included in the analysis. Mean age (±SD) across the entire study population was 44.7±15.7 years and mean symptom duration across the entire study population (±SD) was 12.1±11.8 years.

| MAPP-II (N=378) |  | Male (N=138) |  | Female (N=240) |  | 2020 US Census |
| --- | --- | --- | --- | --- | --- | --- |
|  |  | (SD) |  | (SD) |  |  |
| Age | (years) | 48.5 | 15.4 | 42.6 | 15.6 |  |
| Symptom Duration | (years) | 10.8 | 10.4 | 12.8 | 12.5 |  |
| Race |  | (N) | (%) | (N) | (%) | (%) |
|  | American Indian | 2 | 1.4% | 1 | 0.4% | 1.3% |
|  | Asian | 0 | 0.0% | 3 | 1.3% | 6.1% |
|  | Black | 6 | 4.3% | 12 | 5% | 13.6% |
|  | Native Hawaiian | 0 | 0.0% | 1 | 0.4% | 0.3% |
|  | White | 124 | 89.9% | 213 | 88.8% | 75.8% |
|  | Multi Race | 3 | 2.2% | 9 | 3.8% | 2.9% |
|  | Other | 3 | 2.2% | 1 | 0.4% |  |
|  | Unknown | 0 | 0% | 0 | 0% |  |
| Ethnicity |  |  |  |  |  |  |
|  | Hispanic | 10 | 7.2% | 16 | 6.8% | 18.9% |
|  | Non-Hispanic | 128 | 92.8% | 223 | 92.7% | 59.3% |

### 3.2 Concordance Across All Edges

We compared the two implementations across all 94,830 edgewise models, independent of significance (Table 2, Table 3, Figure 1). Standardized B weights correlated at r = 0.9996 for pain at time of scan and r = 0.9993 for recalled pain. The corresponding test statistics, Satterthwaite-adjusted t for fitlme and Wald z for FEMA, correlated at r = 0.9995 and r = 0.9993 (Figure 1A, 1D), and signed −log10 p values at r = 0.9990 and r = 0.9989. For each of these quantities CCC equaled the Pearson correlation to four decimal places (B, recalled pain: r = 0.9993, CCC = 0.9993), indicating that the small discrepancies between implementations reflect random scatter rather than systematic bias. The median difference in B (FEMA-fitlme) was +0.00004 in both models, with 95% limits of agreement of [−0.0016, +0.0019] and [−0.0017, +0.0020] respectively (Table 3, Figure 1B, 1E). The largest absolute difference in B anywhere in the brain was 0.0095 (pain at time of scan) and 0.0112 (recalled pain), against typical effect magnitudes of 0.11 to 0.13. Marginal R² was highly similar between the implementations (r = 0.9999, CCC = 0.9999 in both models).

**Figure 1.**
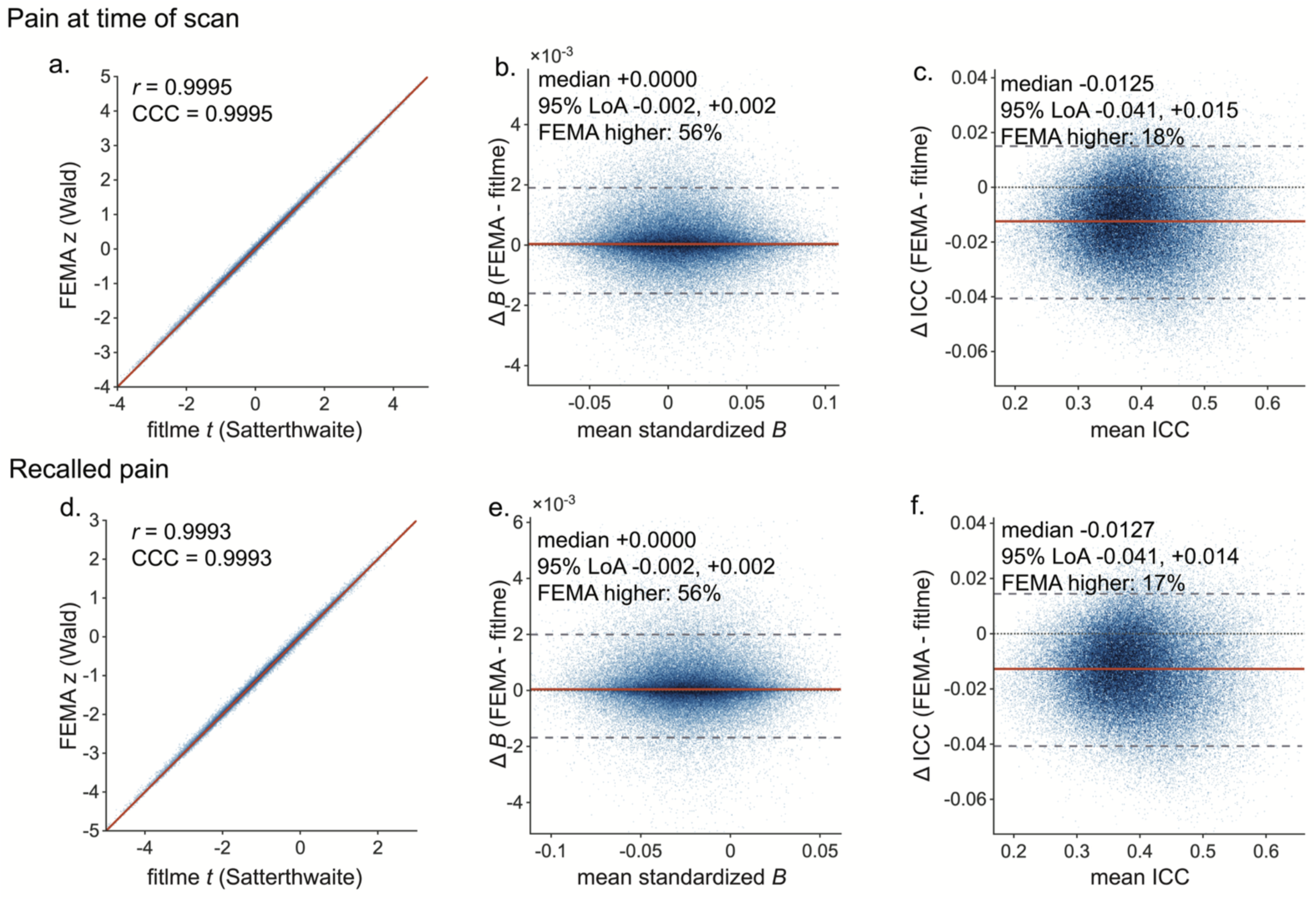
Agreement between the fitlme and FEMA implementations across all 94,830 edgewise models. Top row, pain at time of scan; bottom row, recalled pain (a, d). Test statistics from the two implementations, plotted against the identity line (red) (b, e). Bland-Altman comparison of the standardized fixed effect of pain: the difference between implementations (FEMA minus fitlme) is plotted against their mean (c, f). Bland-Altman comparison of the intraclass correlation coefficient. In panels b, c, e and f the solid orange line marks the median difference, dashed grey lines the 95% limits of agreement (mean ± 1.96 SD), and the dotted line zero.

**Table 2.** Correlation between implementations. Pearson r indexes precision; Lin’s concordance correlation coefficient (CCC) is included for its sensitivity to systematic differences in location or scale. Test statistics are Satterthwaite-adjusted t for fitlme and Wald z for FEMA, each computed from that implementation’s own standard error.

|  | Pain at time of scan |  | Recalled pain |  |
| --- | --- | --- | --- | --- |
|  | Pearson r | CCC | Pearson r | CCC |
| Standardized $\beta$ (fixed effect of pain) | 0.9996 | 0.9996 | 0.9993 | 0.9993 |
| Test statistic (t / z) | 0.9995 | 0.9995 | 0.9993 | 0.9993 |
| Signed $-\log_{10} p$ | 0.9990 | 0.9990 | 0.9989 | 0.9989 |
| Variance explained by fixed effects (marginal $R^2$) | 0.9999 | 0.9999 | 0.9999 | 0.9999 |
| Residual variance | 0.9926 | 0.9878 | 0.9927 | 0.9877 |
| Standard error of $\beta$ | 0.9867 | 0.9852 | 0.9865 | 0.9851 |
| Variance explained by full model (conditional $R^2$) | 0.9862 | 0.9753 | 0.9865 | 0.9751 |
| Intraclass correlation coefficient (ICC) | 0.9826 | 0.9689 | 0.9830 | 0.9686 |
| Random-effect (participant) variance | 0.9745 | 0.9525 | 0.9750 | 0.9520 |

**Table 3.** Difference between implementations (FEMA minus fitlme). Distribution of the edgewise difference between implementations. LoA, limits of agreement (mean difference ± 1.96 SD).

|  | Pain at time of scan |  | Recalled pain |  |
| --- | --- | --- | --- | --- |
|  | Median (Q1, Q3) | 95% LoA | Median (Q1, Q3) | 95% LoA |
| Standardized $\beta$ | +0.00004 (−0.00022, +0.00045) | −0.0016, +0.0019 | +0.00004 (−0.00022, +0.00047) | −0.0017, +0.0020 |
| Standard error of $\beta$ | −0.000025 (−0.000095, +0.000063) | −0.00029, +0.00029 | −0.000017 (−0.000088, +0.000078) | −0.00029, +0.00031 |
| ICC | −0.0125 (−0.0220, −0.0032) | −0.0406, +0.0150 | −0.0127 (−0.0223, −0.0037) | −0.0407, +0.0144 |
| Random-effect variance | −0.0146 (−0.0257, −0.0042) | −0.0477, +0.0169 | −0.0150 (−0.0260, −0.0047) | −0.0477, +0.0163 |
| Residual variance | +0.0076 (+0.0012, +0.0143) | −0.0115, +0.0273 | +0.0079 (+0.0015, +0.0145) | −0.0110, +0.0274 |

Agreement in the estimated variance components was weaker and directionally biased. Intraclass correlation coefficients correlated at r = 0.983 in both models, but CCC was lower (0.969 and 0.969). Random-effect variance agreement was also lower (r = 0.975 and 0.975; CCC = 0.953 and 0.952). The gap between the two coefficients indicates a consistent offset. FEMA attributed less variance to the participant random effect than REML (median difference −0.0146 and −0.0150) and more to residual error (median difference +0.0076 and +0.0079), yielding a lower ICC in approximately 82% of edges in both models (median difference −0.0125 and −0.0127). Bland-Altman comparison (Figure 1C, 1F) showed this offset was uniform across the range of estimated ICC values.

### 3.3 Concordance Across Significant Edges

Both FEMA and the fitlme approach identified connectivity within a sparse set of sensory-discriminative regions as being associated with pain at time of scan, while past-week recalled pain was associated with distributed patterns across regions implicated in memory, salience, and affective processing (Figure 2).

**Figure 2.**
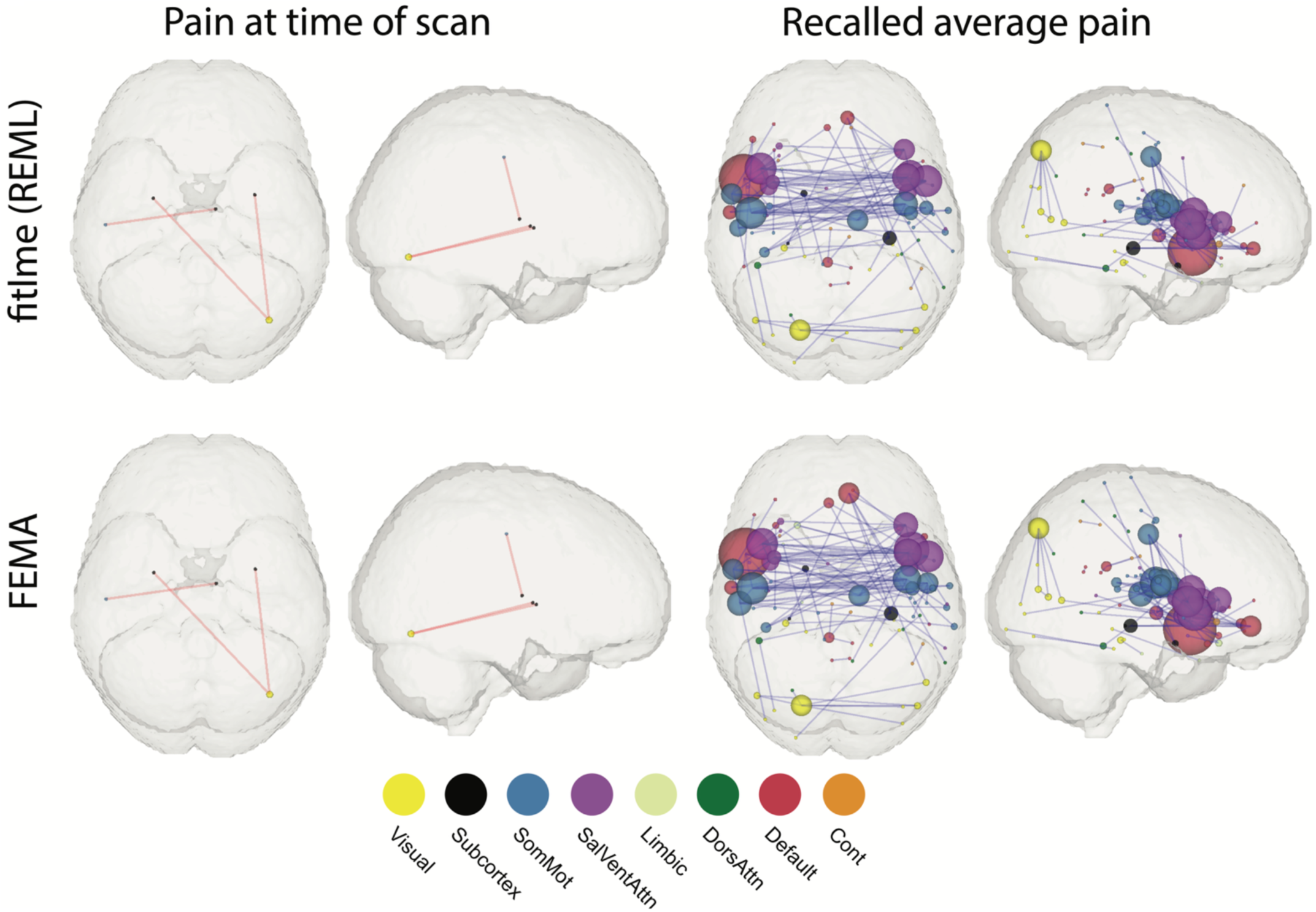
Comparison of significant edges identified in the pain at time of scan and recalled pain analyses for the two approaches (fitlme vs FEMA). Vis=Visual, SomMot=somatosensory/motor, SalVentAttn=Salience Ventral Attention, Cont=Executive Control. Spheres at each node are scaled by the number of significant edges they belong to. Spheres are color coded based on the network they belong to in the Schaefer/Brainnetome atlas. Line color indicates positive (red) or negative (blue) beta weight.

For the pain at time of scan analysis, fitlme and FEMA both produced three, overlapping significant edges. The three edges implicated in the pain at time of scan analysis were the right occipital fusiform gyrus to bilateral basal ganglia and left postcentral gyrus to right thalamus, involving a total of five unique nodes. For the recalled pain analysis, fitlme produced 104 significant edges, and FEMA produced 113. 99 of these edges were overlapping between the two methods. There were 5 edges unique to the fitlme approach and 14 unique to FEMA. Set-level agreement was complete for pain at time of scan (Dice = 1.00) and high for recalled pain (Dice = 0.912; Jaccard = 0.839); treating the fitlme implementation as the reference, FEMA recovered 95.2% of significant edges, and the two implementations reached the same conclusion for 99.98% of the 94,830 edges tested (Supplementary Table 3). Summary statistics for overlapping significant edges can be found in Table 4. Model information for non-overlapping edges from the recalled pain analysis can be found in Supplementary Table 1.

**Table 4.** Model statistics for concordantly significant edges. Model statistics summarized across edges reaching significance under both implementations (q < 0.01): three edges for pain at time of scan and 99 for recalled pain. Because only three edges were concordantly significant in pain at time of scan analysis, values for that model are reported as median (range). Recalled pain values are median (Q1, Q3).

|  | Pain at time of scan<br>median (range), n = 3 |  | Recalled pain<br>median (Q1, Q3), n = 99 |  |
| --- | --- | --- | --- | --- |
|  | fitlme | FEMA | fitlme | FEMA |
| Standardized $\beta$, fixed effect of pain | 0.1321<br>(0.1256 - 0.1356) | 0.1326<br>(0.1257 - 0.1355) | -0.1117<br>(-0.1162, -0.1080) | -0.1112<br>(-0.1162, -0.1080) |
| Standard error of $\beta$ | 0.0252<br>(0.0242 - 0.0252) | 0.0250<br>(0.0238 - 0.0253) | 0.0236 (0.0233, 0.0242) | 0.0237 (0.0235, 0.0242) |
| Intraclass correlation coefficient (ICC) | 0.3884<br>(0.3802 - 0.4452) | 0.4055<br>(0.3959 - 0.4517) | 0.4611<br>(0.4148, 0.4881) | 0.4358<br>(0.4021, 0.4701) |
| Variance explained by fixed effects (marginal $R^2$) | 0.0229<br>(0.0222 - 0.0671) | 0.0228<br>(0.0222 - 0.0669) | 0.0381<br>(0.0273, 0.0560) | 0.0383<br>(0.0275, 0.0572) |
| Variance explained by full model (conditional $R^2$) | 0.4020<br>(0.3944 - 0.4824) | 0.4187<br>(0.4097 - 0.4884) | 0.4909<br>(0.4403, 0.5130) | 0.4685<br>(0.4246, 0.4971) |

We additionally examined the unique edges from the recalled pain analysis in the set of model outputs from the opposite model approach: that is, for edges that were only significant in the fitlme analysis, we examined the model statistics for that same edge in the FEMA analysis. We did the same for the edges uniquely significant in the FEMA analysis. The significant connections unique to FEMA were all edges in which models successfully converged in the fitlme analysis. Upon inspection of the model statistics for these edges in fitlme, all 14 edges were approaching significance, with q values between 0.0100 and 0.0118. The same was true of the 5 edges unique to the fitlme analysis. Model statistics for these non-overlapping edges have been included as Supplementary Table 1.

Overlapping significant edges were highly similar for the fixed effects of pain. FEMA and fitlme agreed on direction of effects for all overlapping edges in both the pain at time of scan and recalled pain analyses. Because only three edges reached significance in the pain at time of scan analysis, correlations computed across this set are uninformative; whole-brain concordance for this model is reported above. For recalled pain, the p values, q values, and normalized B weights across the 99 concordantly significant edges were also highly correlated (r = 0.933, 0.956, and 0.996, respectively), although these values are attenuated by range restriction relative to the whole-brain estimates.

In addition to arriving at a similar set of findings, FEMA offered a reduction in processing time: FEMA completed each whole-brain analysis in 21-35 seconds, compared with 8-10 minutes (473-602 s) for a fitlme implementation parallelized across 14 workers. With no parallel processing, the fitlme approach took approximately 65 minutes per whole-brain analysis.

### 3.4 Sensitivity to FEMA’s Variance-Grid Resolution

Estimated variance components were numerically identical at every grid resolution tested but standard errors changed. Agreement with the fitlme implementation in the estimated standard error improved from r = 0.987 at the default resolution (20 bins) to r = 0.996 at 100 bins, with negligible further change at 1000 bins and no binning.

For recalled pain, FEMA identified 113 significant edges at the default resolution and 100 at all finer resolutions, against the fitlme reference of 104. The number of discordant edges fell from 19 (5 unique to fitlme, 14 unique to FEMA) to 8 (6 and 2, respectively), and the Dice coefficient rose from 0.912 to 0.961. Twelve of the 14 edges uniquely identified by FEMA at the default resolution were therefore attributable to grid discretization rather than to a difference in model behavior. For pain at time of scan, the same three edges were identified at every resolution.

Computation time increased from 21-35 seconds at the default resolution to 40-58 seconds at 100 bins and 89-101 seconds with binning disabled, all of which remained substantially faster than the parallelized fitlme implementation. Full results are given in Supplementary Table 2.

## 4 Discussion

Here we apply FEMA (Fast and Efficient Mixed-Effects Algorithm for large-sample whole-brain imaging data) to a non-ABCD dataset to analyze whole-brain functional connectivity in individuals with chronic pain. Across two complementary dimensions of the pain experience (pain reported at the time of scanning and recalled average pain over the preceding week) FEMA produced results that were highly concordant with those obtained using a likelihood-based mixed-effects solver. In addition to producing highly similar results, FEMA offered a substantial reduction in computation time. Whole-brain analyses that required several minutes using a parallelized fitlme implementation were completed in seconds using FEMA. These findings extend the original validation of FEMA (Parekh et al., 2024) by demonstrating its reliability in a non-ABCD dataset, in a clinical population characterized by substantial within-subject variability, and in the context of connectome-wide functional connectivity analyses. To date, FEMA has been applied to neuroimaging data in one other, non-ABCD dataset: the AHEAD study (Bayat et al., 2025; Curtis et al., 2025) based out of Florida International University in Miami, FL. Both papers implemented ABCD compliant preprocessing and processing approaches, and FEMA was used to analyze diffusion weighted imaging (DWI) magnetic resonance imaging (Curtis et al., 2025) and BOLD response to Kiddie-Continuous Performance Test (Bayat et al., 2025).

The neurobiological findings were not introduced as novel patterns here, but rather serve as a consistency check across modeling approaches in a dataset where these effects have been previously reported using standard mixed-effects methods (McLain et al., 2026). Briefly, both approaches identified sparse and overlapping patterns of functional connectivity associated with pain at the time of scan, primarily involving regions implicated in sensory-discriminative processing. In contrast, recalled pain over the preceding week was associated with a more distributed pattern of connectivity spanning regions commonly linked to affective, salience, and memory-related processes. This dissociation aligns with prior conceptualizations of pain as a multidimensional experience, with momentary pain more closely tied to sensory processing (Borsook et al., 2013; De Ridder et al., 2022; Liu & Kelliher, 2022) and recalled or clinical pain reflecting higher-order integrative systems (Barroso et al., 2021; Jefferson et al., 2021; Neugebauer, 2020; Stegemann et al., 2023; Waisman & Katz, 2024).

Model statistics for the fixed effects of pain (normalized *B* weight, p, and q values) were almost perfectly correlated between the two approaches. The significant findings in the pain at time of scan analysis were identical in terms of identified edges (Figure 2) and highly correlated across various model statistics (Table 4). There were some minor inconsistencies in the larger set of recalled pain findings. Out of 113 significant edges from FEMA and 104 edges from fitlme, 99 were overlapping (Table 4). This left 5 edges unique to the fitlme analysis and 14 edges unique to the FEMA analysis (Supplementary Table 1). Inspection of these non-overlapping edges revealed that they were uniformly near the significance threshold in both implementations, suggesting that these differences likely reflect marginal variation in standard error estimation rather than meaningful divergence in model behavior. Several methodological differences between FEMA and conventional mixed-effects solvers may contribute to these small discrepancies.

FEMA and fitlme rely on fundamentally different estimation strategies, with fitlme using restricted maximum likelihood (REML) estimation and FEMA relying on a method-of-moments approach to estimate variance components, paired with generalized least squares for fixed-effect estimation. This is one of the core differences that allows FEMA to complete a similar set of analyses in magnitudes of order shorter time (Parekh et al., 2024). These differences in approach have the potential to produce different standard errors and variance estimates. Moment-based estimators are consistent but generally less efficient than REML in finite samples and can become biased when variance estimates are constrained to be non-negative (Swallow & Monahan, 1984). Fixed-effect point estimates, however, are generally robust to such differences, even when the estimated variance components diverge modestly (Kackar & Harville, 1984; Kenward & Roger, 1997; Parekh et al., 2024). Consistent with this expectation, we observed near-identical fixed-effect estimates across methods, alongside a small but systematic difference in the variance components: FEMA attributed less variance to the participant random effect and correspondingly more to residual error, yielding a lower ICC in approximately 82% of edges in both models.

Another potential source for the nonoverlapping findings in the recalled pain analysis are the differences in p value determination. The specific fitlme implementation described in this analysis relied on Satterthwaite degrees of freedom to estimate the final p values from fixed effects when derived from a mixed effects structure. The Satterthwaite p-values come from an approximate t/F distribution (Satterthwaite, 1946). By contrast, FEMA treats fixed-effect test statistics as Wald z-scores derived from the standard normal distribution. With moderate/large samples, Satterthwaite t and normal z are very similar (Luke, 2017), which is reflected in the p and q values identified in this analysis. Given the high correlation of p, q, and fixed effects for pain across the two approaches, it seems more likely that the factor driving the difference is related to the calculation of standard error discussed in the previous paragraph.

A secondary analysis additionally indicated that the small number of edges on which the two implementations disagreed was largely attributable to FEMA’s variance-grid approximation rather than to the estimator itself. Increasing the grid resolution from the default of 20 bins to 100 eliminated 12 of the 14 edges uniquely identified by FEMA, and reduced overall disagreement from 19 to 8 of 94,830 edges at a runtime cost of under a minute. Investigators whose conclusions depend on edges close to the correction threshold may therefore wish to increase this bin setting, although the default appears adequate for the type of connectome-wide analysis used in the present analysis.

Notably, the systematic difference in estimated variance components was unaffected by grid resolution, indicating that it reflects a difference between moment-based and likelihood-based estimation rather than an artifact of the approximation.

Another distinction between the two approaches concerns model convergence. Conventional mixed-effects solvers require explicit convergence checks while the FEMA approach does not. Of the 94,830 models per pain measure, none failed to converge under the fitlme implementation, but 183 (pain at time of scan) and 164 (recalled pain) returned an exit status indicating that optimality could not be certified. However, none of these reached significance. Nonconvergence therefore cannot be a driver of the non-overlapping results.

Several limitations of the present work should be noted. While the present dataset includes repeated measures, the random-effects structure was limited to a random intercept for participant. In a previous version of the fitlme implementation, it was discovered that adding a random slope led to overfitting and poor convergence across the 94,830 models. More complex random-effects specifications may reveal additional differences between estimation approaches. Checks for overfitting should always be employed when applying any mixed effects approach, FEMA or otherwise.

In summary, these results demonstrate that FEMA produces results that closely match those obtained using conventional mixed-effects modeling while dramatically improving computational efficiency. By benchmarking FEMA in a longitudinal chronic pain dataset and in a connectome-wide functional connectivity analysis, this work supports its use as a scalable and reliable alternative to traditional mixed-effects solvers. More broadly, FEMA lowers the practical threshold for incorporating mixed-effects modeling into whole-brain neuroimaging analyses, facilitating more rigorous treatment of within-subject variability in studies of complex clinical phenotypes.

## Data and Code Availability

All MAPP neuroimaging data and associated clinical information are available through the NIDDK Central Repository (https://repository.niddk.nih.gov/). Code used to process, analyze, and generate figures from the data is available from the corresponding author on request, and publicly on publication. Code for the FEMA implementation was made available by its authors on GitHub at https://github.com/cmig-research-group/cmig_tools.

## Author Contributions

Natalie McLain: Conceptualization, Methodology, Software, Formal analysis, Investigation, Visualization, Writing - original draft. Chelsea Kaplan: Methodology, Writing - review & editing. Steven Harte: Methodology, Writing - review & editing. Andrew Schrepf: Conceptualization, Methodology, Supervision, Funding acquisition, Writing - review & editing.

## Acknowledgements

The authors thank all of the volunteers who participated in the Symptom Patterns Study. The authors additionally thank the team who developed and made the FEMA algorithm available for public use.

## Supplementary Material

Supplementary Table 1: model statistics for non-overlapping significant edges in the recalled pain analysis. Supplementary Table 2: full results of the variance-grid resolution sensitivity analysis. Supplementary Table 3: edgewise concordance of statistical conclusions across all 94,830 edges.

